# Silencing branched-chain ketoacid dehydrogenase or treatment with branched-chain ketoacids ex vivo inhibits muscle insulin signaling

**DOI:** 10.1101/2020.02.21.960153

**Authors:** Dipsikha Biswas, Khoi T. Dao, Angella Mercer, Andrew Cowie, Luke Duffley, Yassine El Hiani, Petra C. Kienesberger, Thomas Pulinilkunnil

## Abstract

Branched-chain α-keto acids (BCKAs) are downstream catabolites of branched-chain amino acids (BCAAs). Mitochondrial oxidation of BCKAs is catalyzed by branched-chain ketoacid dehydrogenase (BCKDH), an enzyme sensitive to inhibitory phosphorylation by BCKD kinase (BCKDK). Emerging studies show that defective BCAA catabolism and elevated BCKAs levels correlate with glucose intolerance and cardiac dysfunction. However, if/how BCKDH and BCKDK exert control on the availability and flux of intramyocellular BCKAs and if BCKA reprograms nutrient metabolism by influencing insulin action remains unexplored. We observed altered BCAA catabolizing enzyme expression in the murine heart and skeletal muscle during physiological fasting and diet-induced obesity and after ex vivo exposure of C2C12 cells to increasing concentration of saturated fatty acid, palmitate. BCKAs per se impaired insulin-induced AKT phosphorylation and AKT activity in skeletal myotubes and cardiomyocytes. In skeletal muscle cells, mTORC1 and protein translation signaling was enhanced by BCKA with concomitant suppression of mitochondrial respiration. Lowering intracellular BCKA levels by genetic and pharmacological activation of BCKDHA enhanced insulin signaling and activated pyruvate dehydrogenase, an effector of glucose oxidation and substrate metabolism. Our findings suggest that BCKAs profoundly influence muscle insulin function, providing new insight into the molecular nexus of BCAA metabolism and signaling with cellular insulin action and respiration.

## 1. Introduction

Branched-chain amino acids (BCAAs) and their catabolites are bioactive molecules with a broad repertoire of metabolic actions in cellular health and disease (1–6). Leucine, isoleucine and valine (BCAAs) are reversibly transaminated by branched-chain aminotransferase (BCAT) to yield branched-chain α-keto acids (BCKA), specifically α-ketoisocaproate or ketoleucine (α-KIC), α-ketoisovalerate or ketovaline (α-KIV), and α-kethomethylvalerate or ketoisoleucine (α-KMV). BCKAs are irreversibly decarboxylated by BCKA dehydrogenase (BCKDH), a rate-limiting step in BCAA catabolism. BCKDH activity is tightly regulated by BCKA dehydrogenase kinase (BCKDK) mediated inhibitory phosphorylation (7–9) of BCKDH or BCKDH dephosphorylation and activation by protein phosphatase 2C (PP2Cm)(4, 10). Transcriptional regulation of the BCAA catabolizing enzymes and BCAA catabolism is controlled by Kruppel like factor 15 (KLF15) (11), which also regulates glucose and fat oxidation (12–14). Emerging studies have shown that not only inborn mutations in BCAA enzymes, but also dysfunctional BCKA oxidation manifest in metabolic disorders like cancer (15, 16), obesity and insulin resistance (17–19) ischemia (20) and, diabetic cardiomyopathy (21, 22).

Skeletal and cardiac muscle insulin signaling and sensitivity are critical for metabolic homeostasis and organ function (23, 24). A strong association between plasma BCAAs, muscle BCAA catabolic gene expression and insulin resistance (IR) has been reported in several human and rodent models of obesity (25, 26). Whether defects in muscle BCAA catabolizing enzyme expression and activity precede or follow obesity-induced IR remains unclear. Strikingly, pharmacological inhibition of BCKDK in ob/ob and diet-induced obese (DIO) mice normalized BCAA catabolism and attenuated IR (26), signifying that restoring BCAA catabolism and clearing BCAA metabolites in endocrine disorders has beneficial functional outcomes. Plausibly a metabolic environment favouring the accumulation of BCAA catabolites reprograms substrate utilization and causes metabolic dysfunction. However, it remains unknown if BCKAs directly impact muscle insulin signaling to influence substrate utilization and mitochondrial function. A recent report demonstrated that incubation of cardiomyocytes with high levels of BCKAs reduces glucose uptake (27) and remodels mitochondrial respiration (28), questioning the effect of BCKA on insulin signaling and action.

In this study, we investigated if acute fasting and chronic diet-induced obesity alter the expression of BCAA catabolic enzymes, and if this effect is attributed to insulin action or lipid overload. We examined the influence of individual BCKAs on both cardiac and skeletal muscle insulin signaling and protein translation machinery including glucose and mitochondrial metabolism. Lastly, we examined whether targeting BCAA catabolizing enzymes was sufficient to alter intracellular BCKA levels to impact muscle insulin signaling.

## 2. Results

### 2.1 Overnight fasting augments BCKDH phosphorylation and protein expression in the cardiac tissue but not in the gastrocnemius muscle

We examined whether acutely limiting nutrient intake (fasting) alters transcriptional or post-translational levels of BCAA catabolic enzymes in the cardiac and skeletal muscle (Fig 1a). Male C57BL/6 mice subjected to overnight fasting exhibited a decline in body weight, serum glucose and liver weight (Table S1) when compared to either ad libitum fed or refed mice. Ventricular weight to body weight ratio remained unchanged between groups (Table S1). mRNA expression of BCAA catabolic enzymes (*Bckdha, Bckdk, Bcat2, Klf15, Oxct2a, Ivd2, Hmgcs1* and *Mut*) was similar between fed, fasted and refed groups in gastrocnemius and cardiac muscle (Fig 1b-c). Inactivating phosphorylation of BCKDE1α at Ser 293 (suggestive of decreased BCKDHA activity) and total protein content of BCKDHA and BCKDK were unaltered in the gastrocnemius muscle (Fig 1d). However, a marked increase in phosphorylation of BCKDE1α at Ser 293 was observed in the fasted hearts, which returned to fed levels upon 4 h of refeeding (Fig 1e). Interestingly, BCKDHA protein levels were decreased upon fasting and were restored in the hearts of refed mice (Fig 1e). In the heart, BCKDK protein levels remained unaltered across all the groups (Fig 1e). Prior studies in the liver have demonstrated that BCKDHA phosphorylation is decreased following refeeding resulting in increased BCAA oxidation (29, 30) and inhibition of fatty acid oxidation and ketone body generation (31). During fasting, reduced insulin relieves inhibition of fatty acid oxidation in the heart. We propose that the inactivation of BCKDHA downregulates BCAA oxidation thereby rendering fatty acids as dominant substrates for oxidation during fasting. Indeed, leucine inhibited palmitate oxidation in aerobically (30 min) perfused mouse heart (Fig 1f). Since muscle BCAA catabolizing enzyme expression decreased in the presence of excess lipids (32) we postulate that BCAA and fatty acids are competing substrates in the heart likely regulating expression and activity of enzymes facilitating their respective catabolism by affecting insulin signaling.

**Fig 1.**
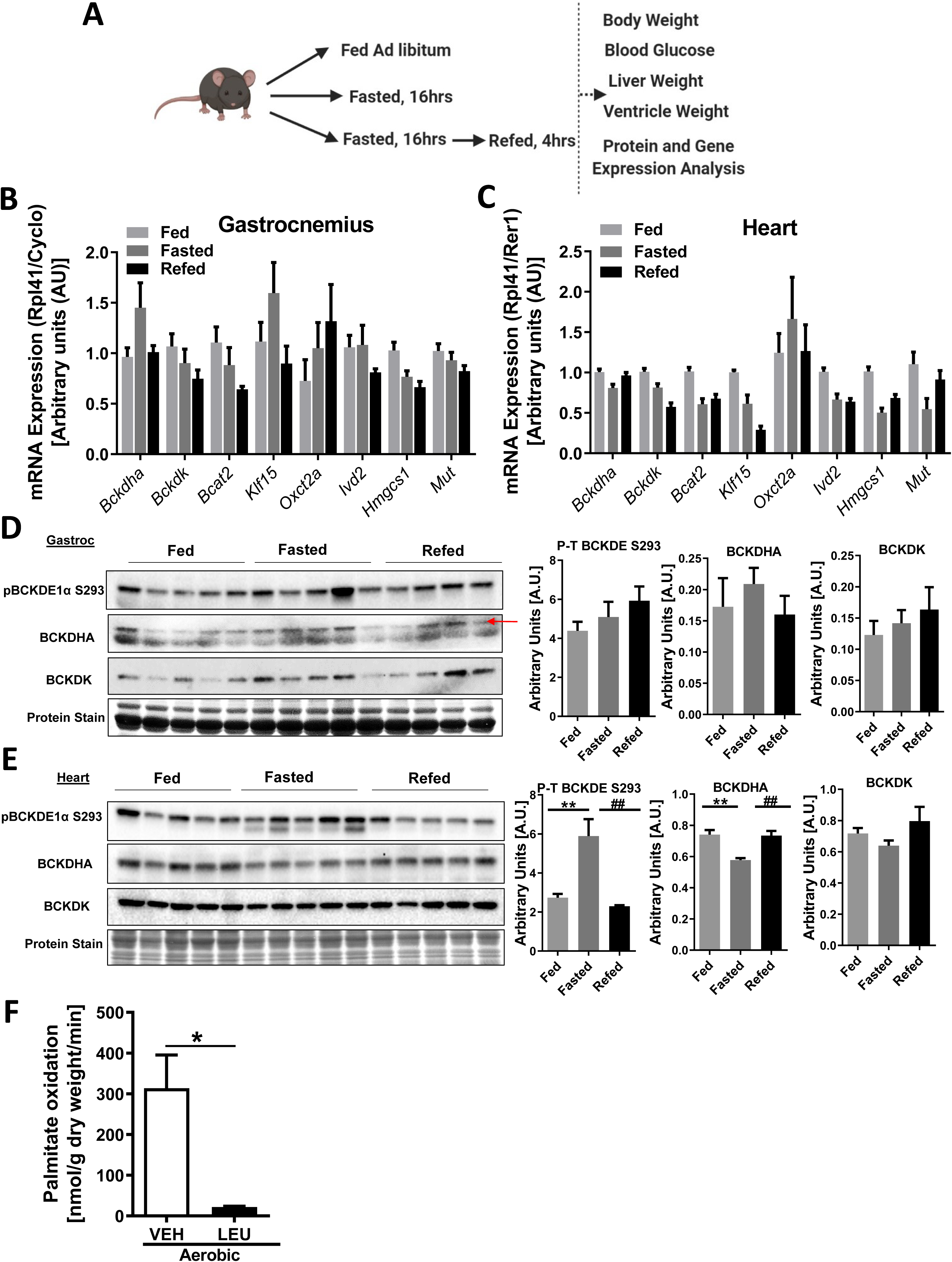
Fasting induces post-translational changes in BCAA catabolic enzymes in the cardiac but not skeletal muscle. a) Study design of C57/BL6 mice ad libitum fed or fasted for 16 hr or refed for 4 hr following fasting, n=5 each group. Quantification of *Bckdha*, *Bckdk*, *Bcat2*, *Klf15*, *Oxct2a*, *Ivd2*, *Hmgcs1*, *Mut* mRNA expression corrected to the reference genes, *Rpl41/Cyclo* in gastrocnemius muscle (b) and *Rpl41/Rer1* in the heart (c). Immunoblot and densitometric analysis of total and phosphorylated Bckdha E1α Ser 293 and total Bckdk in the gastrocnemius muscle (d) and heart (e). Statistical analysis was performed using a two-way ANOVA followed by a Tukey’s multiple comparison test; *p <0,05, **p < 0.01, **** p <0.0001 as indicated. f) Palmitate oxidation rate in isolated mice heart (n=3 per group) perfused aerobically for 30 min with 5mM leucine in the presence of 0.8mM palmitate bound to 2% fatty acid-free bovine serum albumin and 5mM glucose. Graph represents mean ± S.E.M., n=3, *p<0.05 was performed using Student’s t-test.

### 2.2 Expression of BCAA catabolic enzymes is differentially regulated in the cardiac and skeletal muscles following DIO

We next ascertained if amino acid metabolizing enzymes in the muscle were temporally regulated in a model of progressive lipid excess and hyperinsulinemia (diet-induced obesity). Male C65/BL6 mice were fed a high-fat high sucrose (HFHS) diet and blood and tissues were collected at 2, 4, 8, and 13 weeks post feeding to determine the BCAA catabolic enzyme expression in the gastrocnemius muscle and heart (Fig 2a). To examine the effect of progressive lipid loading on BCAA metabolizing enzymes, we compared the expression profile of genes involved in BCAA catabolism in the gastrocnemius muscles between 2wk and 13 wk of HFHS feeding. mRNA expression of *Mut* and *Ivd2*, intermediary effector enzymes of the BCAA oxidation pathway, was reduced in the gastrocnemius muscle at 13 wk compared to 2wk HFHS fed mice (Fig 2b), consistent with a prior report in mice fed a (60%) high fat diet (HFD) (32). Interestingly, *Hmgcs1* levels were upregulated in the gastrocnemius muscle in 13 wk fed mice compared to 2 wk fed mice (Fig 2b), possibly as a compensatory mechanism to maintain CoA pools. In the heart, mRNA expression of the BCAA catabolic enzymes, except for KLF15, remained unaltered in a setting of DIO (Fig 2c). Since the earliest changes in mRNA expression were not detected until 13 wk, we used 4 wk, and not 2 wk, HFHS-fed mice as controls to analyze protein expression. Phosphorylation of BCKDE1α at Ser 293 in the gastrocnemius muscles was unchanged at 8 wk but increased at 13 wk in HFHS-fed mice compared to the 4 wk HFHS-fed mice (Fig 2d). Unlike the fasted state (acute lipid turnover), no changes in BCKDH complex phosphorylation was found in hearts from HFHS fed mice (chronic lipid excess) compared to chow (Fig 2e). Together, these data demonstrate that unlike acute fasting, chronic HFHS feeding downregulates enzymes involved in BCAA catabolism in the gastrocnemius muscle but not the heart.

**Fig 2.**
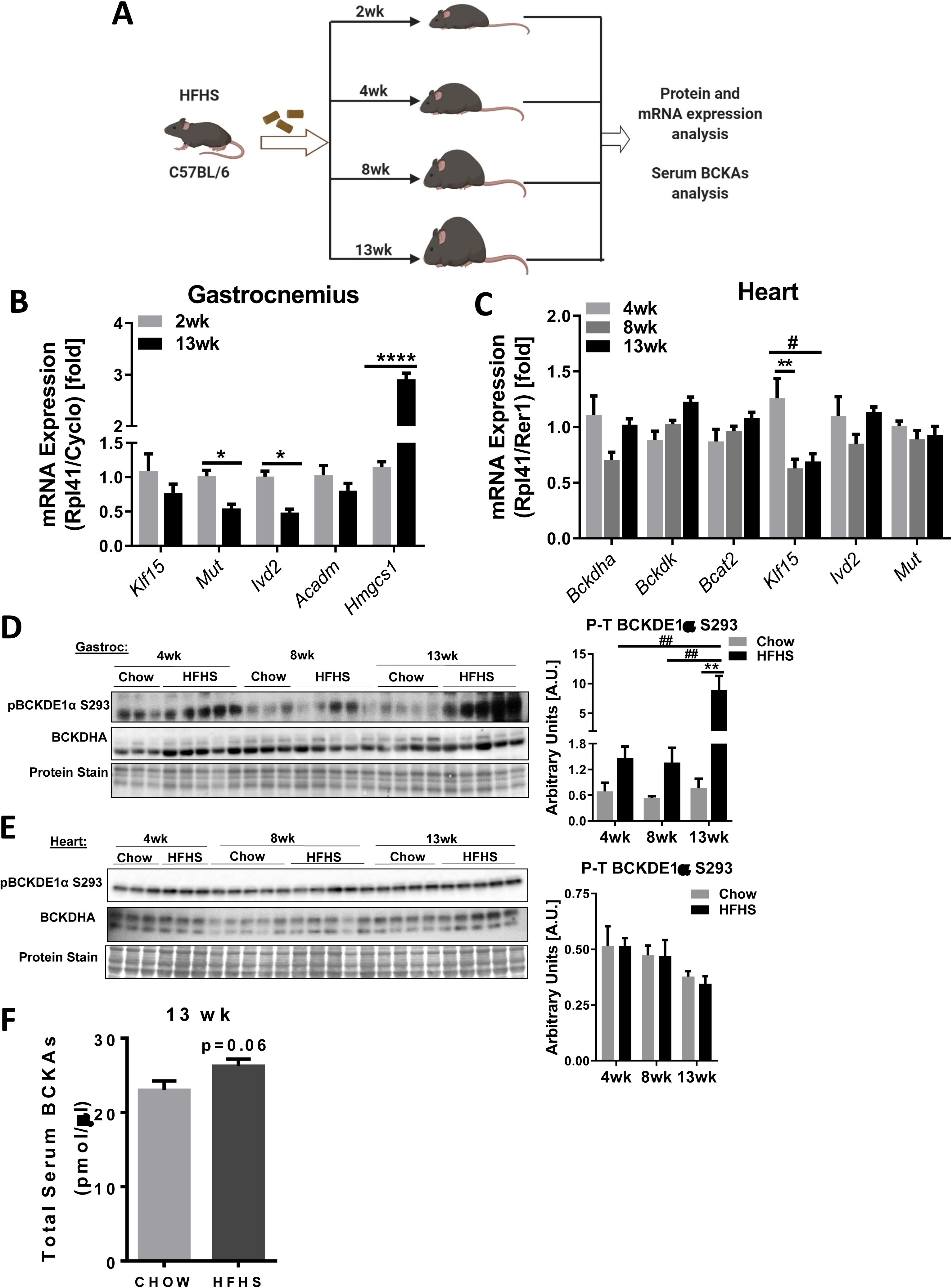
Diet-induced obesity downregulates BCAA catabolizing enzymes in the skeletal muscle. a) Study design of HFHS diet feeding of C57BL/6 mice at 2, 4, 8 and 13 wk. b) Quantification of *Klf15*, *Mut*, *Ivd2*, *Acadm* and *Hmgcs1* mRNA expression corrected to *Rpl41/Cyclo* reference genes in gastrocnemius muscle of 2 wk and 13 wk HFHS fed mice. (c) Quantification of *Bckdha*, *Bckdk*, *Bcat2*, *Klf15*, *Ivd2* and *Mut* mRNA levels corrected to *Rpl41/Rer1* reference genes in the heart of 4 wk, 8 wk and 13 wk HFHS fed mice. (d-e) Immunoblot and densitometric analysis of total and phosphorylated and total Bckdha E1α Ser 293 in gastrocnemius muscle (d) and heart (e). f) Serum BCKAs in 13 wk chow and HFHS diet-fed mice, measured using UPLC MSMS (n=5). Statistical analysis was performed using a two-way ANOVA followed by a Tukey’s multiple comparison test; *p <0,05, **p < 0.01, **** p <0.0001 as indicated.

Chronic lipid overload not only drives IR but also increases systemic and myocellular BCAAs (11, 25) and BCKAs (11). To simulate the gradual increase in fatty acids observed in obesity and IR *ex vivo*, C2C12 cells were treated with low (0.2 mM) and high (0.4mM) palmitate for 16 h. A marked downregulation of *Ivd2* mRNA levels with concomitant upregulation of *Hmgcs1* was observed in C2C12 cells (Fig S1a) consistent with the DIO data (Fig 2b). Since lipid overload altered the expression of distinct BCAA catabolic enzymes, at least in the skeletal muscle, we raised the question if catabolic intermediates of BCAA metabolism, such as BCKAs, influence muscle insulin signaling. Furthermore, we observed a trend towards increased total fasted serum BCKAs as early as 13 wk of HFHS diet feeding (Fig 2f), suggesting that impaired BCAA catabolism in obesity not only elevates levels of BCAAs but also augments BCKAs (17, 33).

### 2.3 BCKA inhibit insulin signalling in skeletal and cardiac muscle cells

BCKAs are BCAA metabolites that are now emerging as clinically relevant biomarkers for IR (34, 35). BCKA are not only generated intracellularly but also transported between tissues to support nitrogen balance, ketone body metabolism, and gluconeogenesis (36, 37). Short term (30 min) treatment of C2C12 cells with increasing concentrations (1 mM, 2 mM and 5 mM) of 4-methyl 2-oxopentanoic acid sodium salt (ketoleucine) resulted in a concentration-dependent impairment of insulin-induced AKT isoform 1 phosphorylation at Ser 473 (Fig S2a). Notably, insulin-mediated activating phosphorylation of insulin receptor substrate 1 (IRS 1) at Tyr 612 was inhibited by ketoleucine at a concentration of 5 mM but not 1 or 2 mM (Fig S2a). Ketoleucine mediated inhibition of AKT1 and IRS1 phosphorylation was observed at 15 min and persisted until 30 min of ketoleucine treatment (Fig S2b). However, phosphorylated AKT1 levels were restored to control levels at 60 min (Fig S2b). The inhibitory effect of ketoleucine on insulin-stimulated AKT1 phosphorylation in C2C12 cells was also recapitulated in the rat L6 muscle cell line (Fig S2c). We next tested if BCKAs modulated palmitate-induced IR. In C2C12 (Fig 3a) and L6 cells (Fig S2d) incubated with 0.4 mM palmitate, ketoleucine did not exacerbate palmitate mediated decreases in AKT1 Ser 473 phosphorylation and phosphorylation of IRS1 at Tyr 612 (Fig 3a). To address whether other ketoacids also impact skeletal muscle insulin signalling, we treated C2C12 cells with 5 mM sodium-3-methyl-2-oxobutyrate (ketovaline) (Fig 3b), 5 mM 3-methyl-2-oxovaleric acid sodium salt (ketoisoleucine) (Fig 3c) and a combination of all BCKA (Fig S2e) in the presence or absence of 0.4 mM palmitate. Similar to the effect of ketoleucine, AKT1 Ser 473 phosphorylation was impaired by both ketovaline and ketoisoleucine in the absence of palmitate (Fig 3b-c). Ketovaline, but not ketoleucine or ketoisoleucine, reduced AKT1 phosphorylation at Thr 308 (Fig 3b). Ketoacids, in combination, significantly reduced AKT1 phosphorylation at Ser 473 in C2C12 cells (Fig S2e). Our data are consistent with a recent report which demonstrated that BCKAs inhibited insulin signalling in 3T3-L1 adipocytes (26). Consistent with reduced AKT1 phosphorylation, insulin induced AKT1 activity was also decreased by ketoleucine and ketovaline (Fig 3d). Ketoisoleucine did not influence AKT1 activity, perhaps reflecting its more modest ability to inhibit insulin-stimulated AKT1 phosphorylation (Fig 3d). Ketoleucine inhibited insulin-mediated AKT1 Ser 473 phosphorylation in the absence of palmitate not only in muscle cells but also in neonatal rat cardiomyocytes (NRCMs) (Fig 4a-b) and adult rat cardiomyocytes (ARCMs) (Fig 4a & c). Similarly, treatment with ketovaline and ketoisoleucine also decreased insulin-stimulated AKT1 S473 phosphorylation in ARCMs (Fig S3a). Taken together, these data suggest that BCKAs per se is equivalent to palmitate in inhibiting insulin signalling in skeletal and cardiac muscle cells.

**Fig 3.**
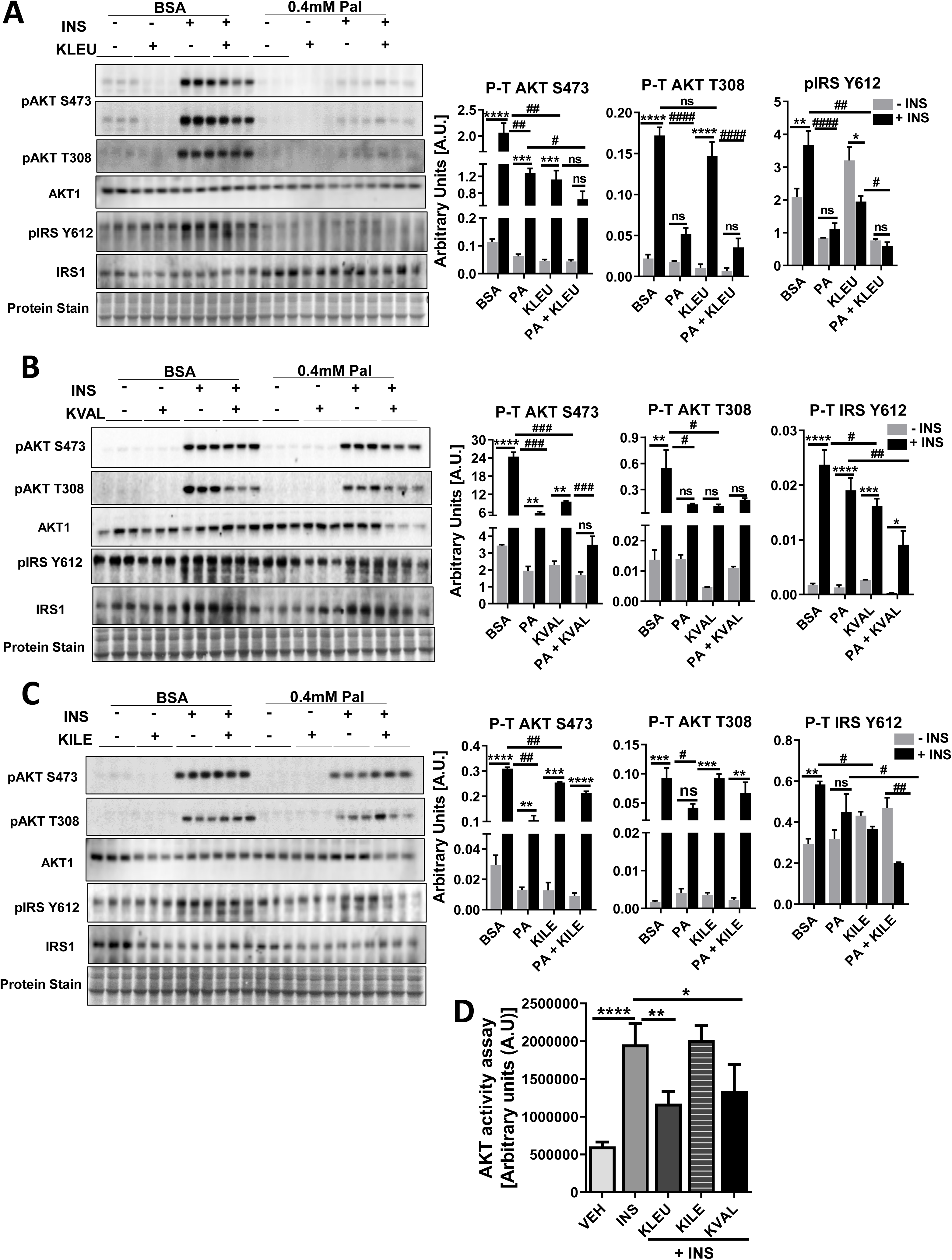
BCKAs impair insulin signaling in C2C12 cells. (a-c) Differentiated C2C12 cells were pre-incubated with either 2% BSA or 2% BSA conjugated with 0.4mM palmitate for 16 hr followed by either a) 5mM ketoleucine, b) ketovaline or c) ketoisoleucine treatment for 30 mins. Myotubes were subjected to 100 nM insulin for 15 mins in presence or absence of individual BCKAs. Immunoblot and densitometric analysis of total and phosphorylated AKT Ser 473 and Thr 309, total and phosphorylated IRS Tyr 612. Statistical analysis was performed using a two-way ANOVA followed by a Tukey’s multiple comparison test; *p <0.05, **p < 0.01, **** p <0.0001 as indicated; *within groups, #between groups. d) Quantification of insulin-induced AKT activity in C2C12 myotubes following 100 nM insulin stimulation for 15mins in the presence or absence of 5mM BCKAs treated for 30 mins. Non-insulin, non-BCKAs treated cells served as controls. The graph represents mean ± S.E.M., n=3, *p<0.05 was performed using Student’s t-test.

**Fig 4.**
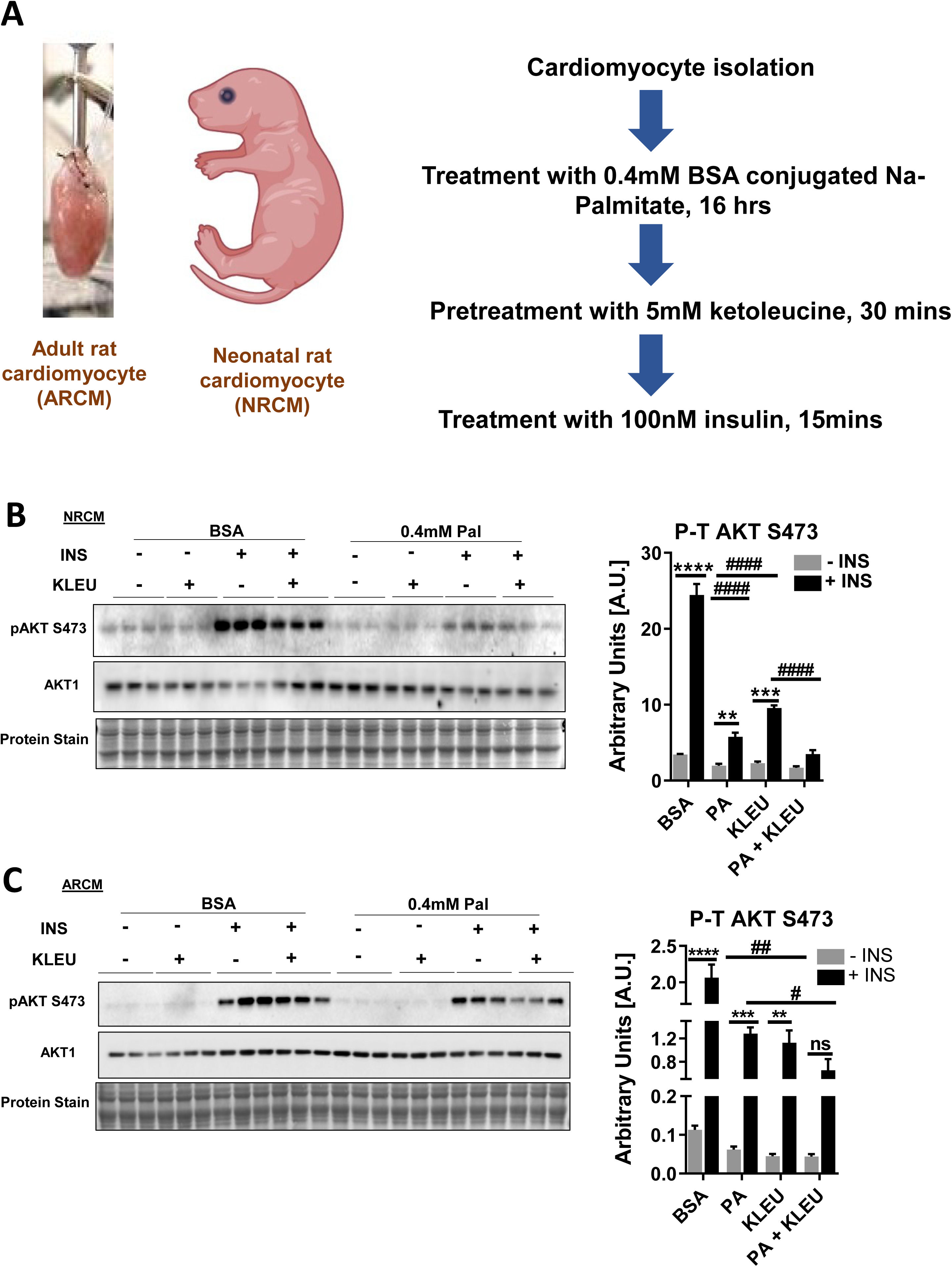
BCKAs impair insulin signaling in cardiomyocytes. a) Schematic of isolation of neonatal (NRCM) and adult (ARCM) rat cardiomyocytes and the treatment protocol. b) NRCMs and c) ARCMs were pre-incubated with either 2% BSA or 2% BSA conjugated with 0.4mM palmitate for 16 hr followed by 5mM ketoleucine for 30 min. Immunoblot and densitometric analysis of total and phosphorylated AKT Ser 473 levels in cardiomyocytes subjected to either 200 nM (NRCMs) or 100 nM (ARCMs) insulin stimulation for 20 mins in the presence or absence of ketoleucine. Statistical analysis was performed using a two-way ANOVA followed by a Tukey’s multiple comparison test; *p <0,05, **p < 0.01, **** p <0.0001 as indicated; *within groups, #between groups.

### 2.4 Ketoleucine blunts insulin-induced decreases in PP2A activity

Since acute BCKAs treatment impaired insulin signaling, we questioned whether BCKA target upstream modulators of AKT phosphorylation. Protein phosphatase 2A (PP2A) dephosphorylates and inactivates AKT (38), whereas insulin inhibits PP2A activity (39, 40), preventing AKT dephosphorylation and increasing AKT activity (40, 41). We examined if ketoleucine alters PP2A activity to modulate insulin signaling (Fig 5a). The presence of ketoleucine prevented the inhibitory effect of insulin on PP2A activity, while pre-treatment with the PP2A inhibitor, okadaic acid (OA), decreased ketoleucine’s effect (Fig 5b). We next examined if changes in PP2A activity corresponded with AKT phosphorylation. Since ketoleucine did not affect AKT Thr 308 phosphorylation, we chose to examine AKT 2 phosphorylation, an additional readout of insulin signaling. OA increased phosphorylation of both AKT isoforms, with a more significant effect observed for AKT1 Ser 473 (Fig 5c). Ketoleucine exhibited a more significant inhibitory effect on AKT2 Ser 474 than AKT Ser 473 phosphorylation (Fig 5c). Pre-treatment with OA partially rescued AKT1 Ser 473 phosphorylation and significantly increased AKT2 Ser 474 phosphorylation (Fig 5c). Interestingly, pre-treatment of cells with ketoleucine followed by OA and insulin treatment augmented AKT phosphorylation more than OA alone (data not shown). These data show that ketoleucine blunts insulin-induced inactivation of PP2A activity.

**Fig 5.**
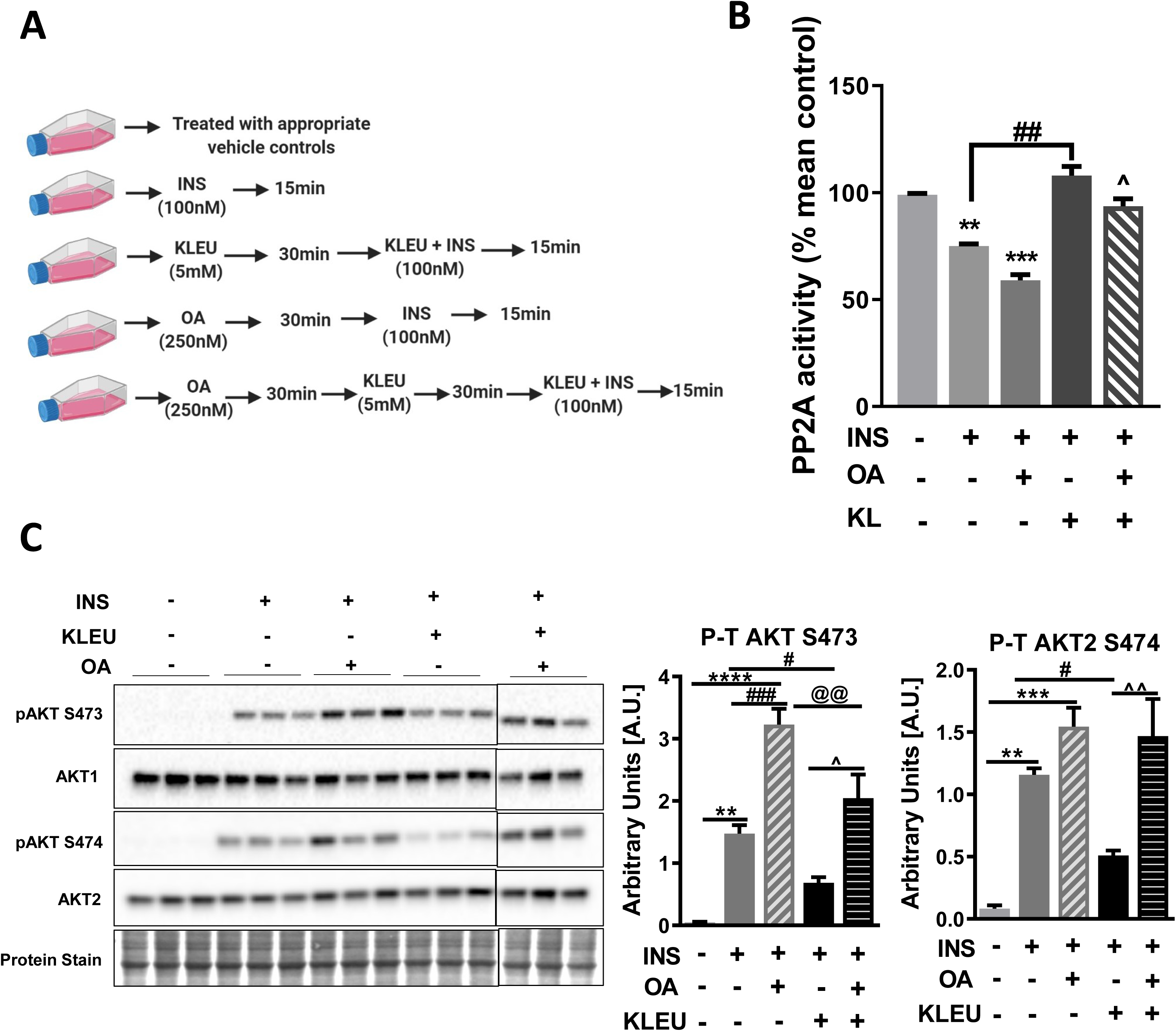
Okadaic acid rescue BCKAs induced insulin signaling impairment. a) Outline of the treatment scheme used to test the effect of okadaic acid on ketoleucine mediated insulin signaling. b) PP2A activity expressed as percent mean control and c) immunoblot and densitometric analysis of total to phosphorylated AKT Ser 473 and total to phosphorylated AKT2 Ser 474 in differentiated C2C12 myotubes pretreated with 250nM okadaic acid (OA) for 30 mins followed by 5mM ketoleucine for 30 mins and 100nM insulin stimulation for 15 mins. No insulin-treated cells were employed as controls. Other relevant groups are provided as an illustration in (a). Statistical analysis was performed using a one-way ANOVA followed by a Tukey’s multiple comparison test; *p <0,05, **p < 0.01, **** p <0.0001 as indicated; *comparison with −INS, #comparison with +INS, ^comparison between +KL+INS v/s OA+KL+INS, @+KL+INS v/s OA+INS.

### 2.5 Affecting clearance of BCKAs by genetic and pharmacological modulation of BCAA catabolizing enzymes influences muscle insulin signalling

We next examined if BCKAs mediated impairment in insulin signaling can be precipitated by increasing endogenous BCKA accumulation as opposed to exposure to exogenous BCKAs. Enzymatic activity of BCKDH determines the catabolic fate of BCKAs (25). BCKDH activity is inhibited by BCKDK-mediated phosphorylation, an effect that is reversed by PPM1K-mediated BCKDH dephosphorylation (Fig 6a). We hypothesized that targeting BCAA catabolizing enzymes will facilitate oxidation and clearance of intracellular BCKAs, rescuing impaired insulin signaling induced by BCKAs. C2C12 myotubes were transduced with adenoviruses overexpressing BCKDK and BCKDHA, the key subunit of BCKDH complex (Fig 6b). Intracellular BCKAs were significantly increased in BCKDK overexpressing cells and decreased in BCKDHA overexpressing cells (Fig 6b), signifying that altering BCAA metabolic enzyme levels changes cellular BCKA content. We next examined if this change in intracellular BCKA levels precipitates alterations in insulin signaling. Consistent with increased accumulation of intracellular BCKAs, BCKDK overexpression decreased insulin-mediated phosphorylation of AKT isoform 1 at Ser 473 and AKT isoform 2 at Ser 474 but not at Thr 308 residue of AKT1 (Fig 6c). AKT1 phosphorylation at both Ser 473 and Thr 308 as well as at the Ser 474 residue of AKT2, were increased in myotubes overexpressing BCKDHA in the presence of insulin (Fig 6d). We observed that BCKDHA was robustly expressed in myoblasts and was undetected by day 4 of differentiation (data not shown). Silencing BCKDHA in C2C12 myoblasts impaired AKT1 Ser 473 phosphorylation, as shown by immunoblot (Fig 6e) and ELISA analysis (Fig S4a). Interestingly, BCKDHA depletion induced a more significant decrease in AKT2 Ser 474 phosphorylation when compared to AKT1 Ser 473 (Fig 6e). Improved insulin signaling induced by BCKDHA overexpression resulted in increased pyruvate dehydrogenase (PDH) activity (Fig S4b). However, BCKDHA silencing did not impact PDH activity (Fig S4b). Similar effects on AKT1 Ser 473 phosphorylation were observed in NRCMs with either overexpression or knockdown of BCKDHA (Fig S4c).

**Fig 6.**
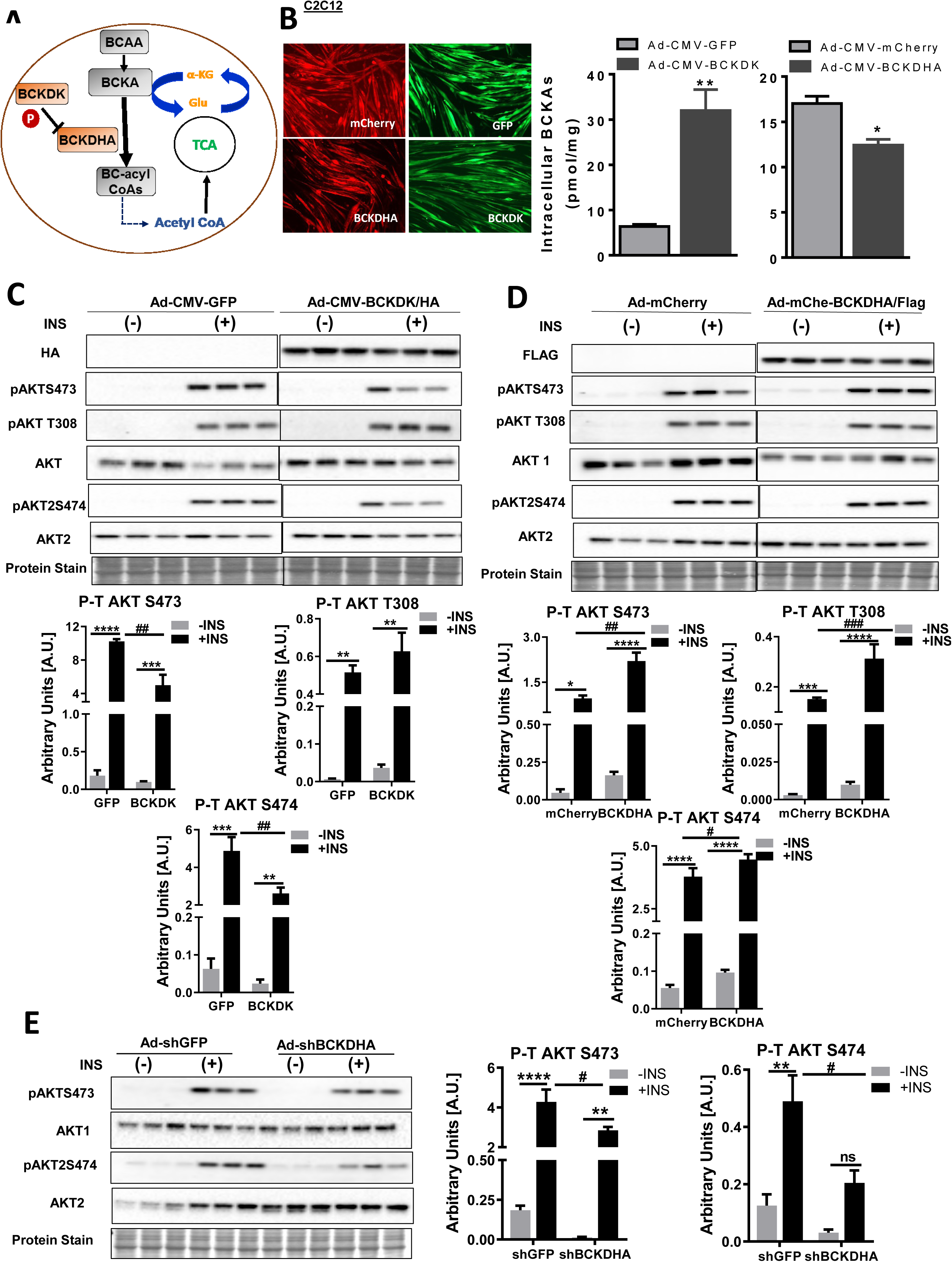
Modulating intracellular BCKAs by modifying BCAA catabolic enzyme expression alters skeletal and cardiac muscle insulin signaling. a) Brief schematic of enzymes regulating BCKA catabolism. b) UPLC mass spectrometric analysis of intracellular BCKAs levels in C2C12 myotubes transduced with Ad-CMV-GFP-hBCKDK/HA (MOI 150) or Ad-CMV-mCherry-hBCKDHA/Flag (MOI 150) and their appropriate controls. The graph represents mean ± S.E.M., n=3, *p<0.05 was performed using Student’s t-test. Immunoblot and densitometric analysis of total and phosphorylated AKT Ser 473, Thr 308 and total and phosphorylated AKT2 Ser 474 in (c) C2C12 myotubes transduced with Ad-CMV-GFP or Ad-CMV-GFP-hBCKDK/HA, (d) Ad-CMV-mCherry or Ad-CMV-mCherry-hBCKDHA/Flag, (e) Ad-CMV-shGFP or Ad-CMV-shGFP-r/mBCKDHA (MOI 200) for 48 hr followed by incubation with 100nM insulin for 15 mins. Statistical analysis was performed using a two-way ANOVA followed by a Tukey’s multiple comparison test; *p <0,05, **p < 0.01, **** p <0.0001 as indicated; *within groups, #between groups.

In addition to genetic manipulation of BCAA catabolizing enzymes, we also employed pharmacological inhibition of BCKDK as an alternative approach to modulate intracellular BCKA levels. 3,6-dichlorobenzothiophene-2-carboxylic acid (BT2) is a selective inhibitor of BCKDK (Fig 7a) which increases BCKDH activity (22, 26) by inhibiting BCKDH phosphorylation. A 20-h exposure of C2C12 myotubes to BT2 reduced intracellular BCKAs (Fig 7b), signifying increased BCKDH activity, consistent with previous reports (22, 26). Furthermore, BT2 markedly increased *Klf15* and moderately increased *Bcat2* mRNA levels (Fig 7c), indicating increased BCKAs flux towards oxidation. BT2 also decreased BCKDE1α Ser 293 inactivating phosphorylation in a concentration-dependent manner (Fig 7d). Further, BT2 increased insulin mediated AKT phosphorylation in C2C12 cells, as shown by immunoblot (Fig 7e) and ELISA analysis (Fig S5a). Similar insulin-dependent effects of BT2 on AKT1 phosphorylation were also observed in NRCMs (Fig S5b). We next questioned whether BT2 could rescue insulin signaling impairment in BCKDHA depleted cells. BT2 restored AKT1 Ser 473 but not AKT2 Ser 474 phosphorylation in BCKDHA silenced C2C12 cells (Fig 7f). BT2 treated C2C12 myotubes displayed increased *Glut1* but not *Glut 4* mRNA levels (Fig 7g). Additionally, BT2 tended to increase insulin-induced pyruvate dehydrogenase (PDH) activity compared to vehicle-treated cells (Fig 7h), supporting the view that BT2 activates PDH to augment muscle glucose oxidation in response to enhanced insulin signaling (42–44). A functional insulin signaling pathway not only signifies a better coupling of glycolysis and glucose oxidation but is an index of efficient lipid oxidation. Indeed, insulin resistance in HFD fed mice is often associated with incomplete fat oxidation and metabolic inflexibility in the skeletal muscle (45, 46). C2C12 cells treated with BT2 displayed increased ^14^C palmitate oxidation (Fig 7i) and a trend towards the reduced accumulation of acid-soluble metabolites (Fig 7j), suggesting that the favourable response of BT2 on insulin signaling is related to BCKA clearance and its downstream effects on lipid oxidation. Our findings are in agreement with a prior study where the key enzyme of fatty acid biosynthesis, ATP citrate lyase, was reported to be inactivated by BT2 (47). Together, these results suggest an intricate integration of BCKAs and lipid metabolism that could be a plausible driver for muscle IR.

**Fig 7.**
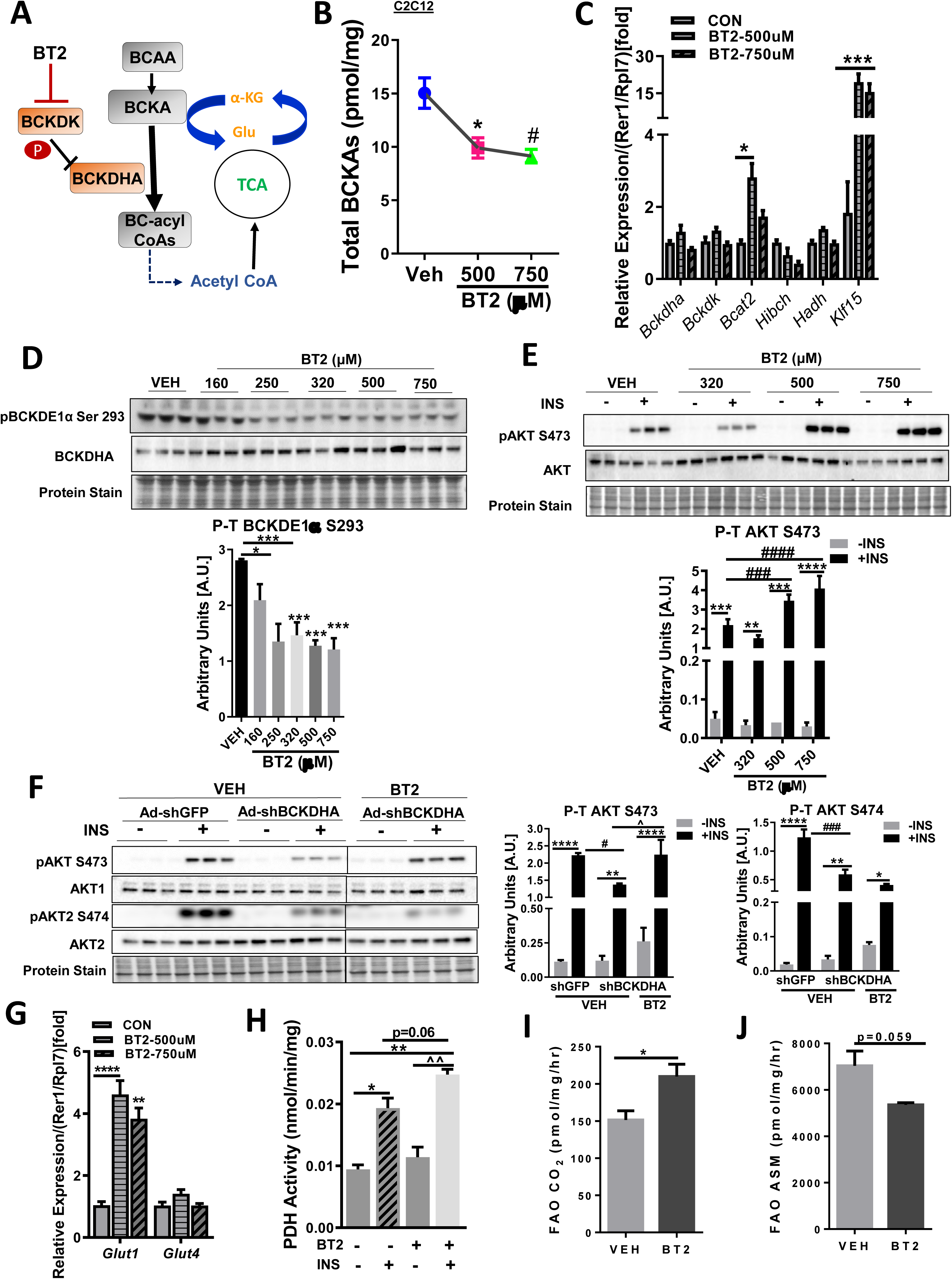
Pharmacological inhibition of BCKDK by BT2 improves muscle insulin signaling. a) A diagrammatic representation of BT2 action. b) Intracellular BCKAs by UPLC/MS-MS and c) qPCR analysis of BCAA catabolic genes *Bckdha*, *Bckdk*, *Bcat2*, *Hadh*, *Hibch* and *Klf15* corrected to Rer1/Rpl7 reference gene levels in C2C12 myotubes treated with 500µM and 750µM BT2 for 20 hr. d) Immunoblot analysis and densitometric quantification of phosphorylated BCKDH subunit E1 at Ser 293 in C2C12 myotubes treated with 160µM, 250µM, 320µM, 500µM and 750µM BT2 for 20 hr. e) Immunoblotting and densitometric quantification of phosphorylated AKT Ser 473 and total AKT in differentiated C2C12 cells pretreated with 320µM, 500µM and 750µM BT2 for 20 hr followed by incubation with 100nM insulin for 15 mins. (f) Immunoblot analysis and densitometric quantification of phosphorylated AKT Ser 473, Ser 474 and total AKT1 and AKT2 in C2C12 myoblasts transduced with Ad-CMV-shGFP or Ad-CMV-shGFP-r/mBCKDHA (MOI 200) for 48 hr followed by incubation with 500µM BT2 for 20 hr and stimulation with 100nM insulin for 15 mins. Statistical analysis was performed using a two-way ANOVA followed by a Tukey’s multiple comparison test; *p <0,05, **p < 0.01, **** p <0.0001 as indicated; *within groups, #comparison with shGFP+INS, ^comparison with shBCKDHA+BT2. g) qPCR analysis of *Glut1* and *Glut4* mRNA levels corrected to Rer1/Rpl7 reference gene levels in C2C12 myotubes treated with or without 500µM BT2 for 20 hr. h) PDH activity analyzed in differentiated C2C12 myotubes preincubated with 500µM BT2 for 20 hr, followed by 100nM insulin treatment for 15mins. Statistical analysis was performed using a one-way ANOVA followed by a Tukey’s multiple comparison test; *p <0,05, **p < 0.01, **** p <0.0001 as indicated; *comparison with −INS, ^comparison with +BT2. ^14^C-palmitate oxidation expressed as i) trapped ^14^CO_2_ or (j) acid-soluble metabolite in C2C12 myotubes treated with or without 500µM BT2 for 20 hr. The graph represents mean ± S.E.M., n=3, *p<0.05 was performed using Student’s t-test.

### 2.6 BCKAs upregulate mTORC1 and protein translation signaling in skeletal muscle cells

BCAAs and BCKAs augment mTOR signaling (48–51) in different cell types (52, 53). We examined if the effect of BCKAs on inhibiting the insulin signaling pathway was associated with increased mTOR signaling (Fig 8a), which is known to negatively regulate insulin signaling (54). Phosphorylation of mTORC1 at Ser 2448 and its downstream target P70S6K at Thr 389 was significantly increased by ketoleucine and ketoisoleucine but not ketovaline (Fig 8b). Next, we assessed the effect of individual BCKAs on other downstream targets of mTOR involved in protein translation. Eukaryotic elongation factor 2 (eEF2) mediates ribosomal translocation during the elongation phase of mRNA translation. Phosphorylation of eEF2 at Thr 56 mediated by eEF2 kinase (eEF2K) interferes with its binding to the ribosome and inhibits translation elongation. Increased mTORC1 signaling inhibits eEF2K and promotes dephosphorylation and activation of eEF2 (55). Consistent with mTORC1 activation, ketoleucine and ketoisoleucine, but not ketovaline, downregulated eEF2 phosphorylation at Thr 56 (Fig 8b). Translation initiation is regulated by mTOR dependent phosphorylation and activation of eukaryotic initiation factor 4G (eIF4G), at Ser 1108 (56). We observed increased eIF4G Ser 1108 phosphorylation in C2C12 cells treated with ketoleucine and ketoisoleucine, but not ketovaline (Fig 8b). Together, our findings are in agreement with a recent study in adipocytes (26) and show that acute incubation with BCKAs, particularly ketoleucine and ketoisoleucine, upregulates mTORC1 activity and protein translation signaling. Consistent with this notion, BCKDHA silencing in C2C12 cells increased phosphorylated mTOR, P70S6K as well as eIF4G (Fig 8c) with concomitant inhibition of insulin signaling (Fig. 6e).

**Fig 8.**
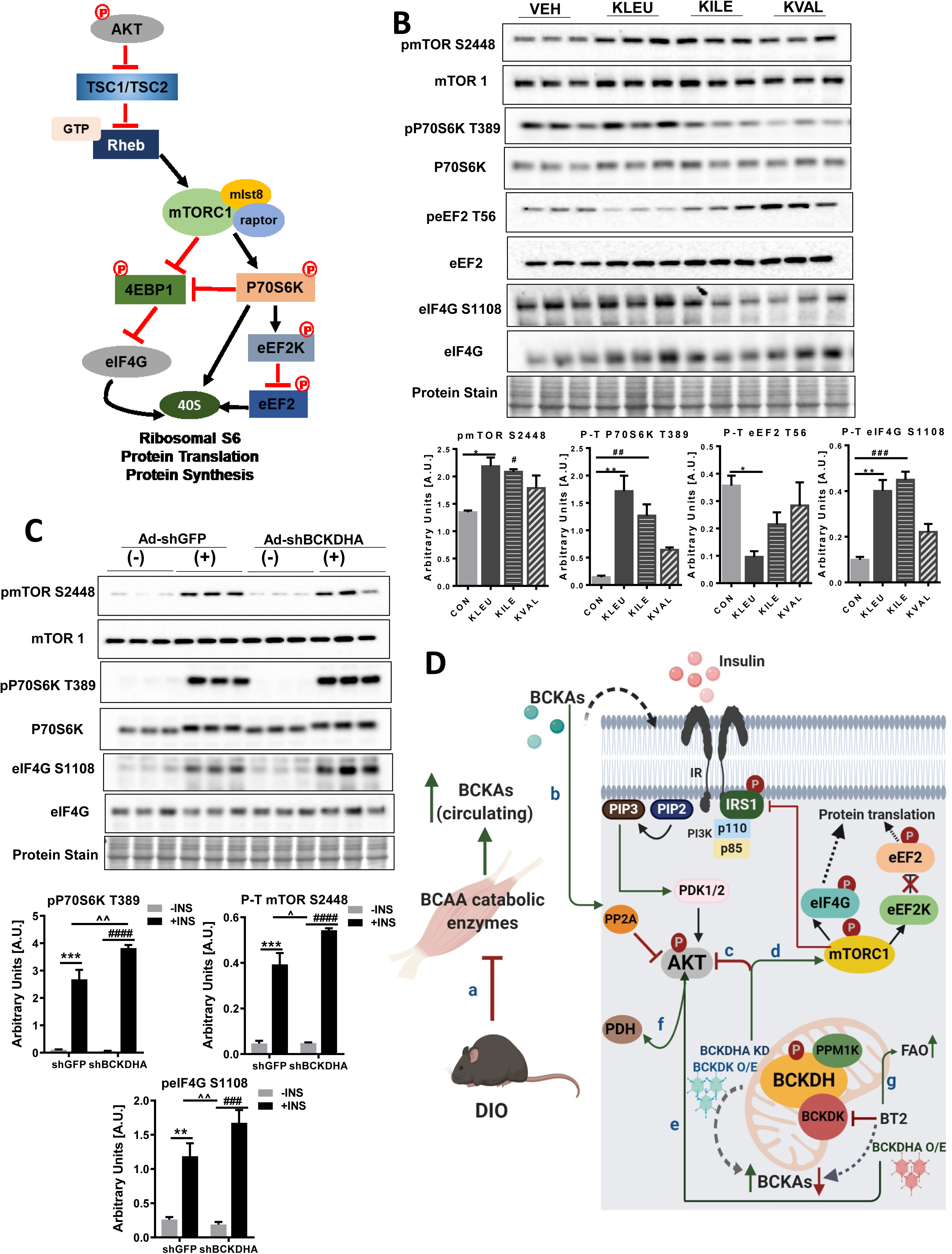
BCKAs activate mTORC1 and downstream protein translation signaling in skeletal muscle. a) Schematic of mTORC1 signaling and its downstream protein translational targets. b) C2C12 myotubes were treated with 5mM ketoleucine, ketoisoleucine and ketovaline for 45 mins. b-c) Immunoblot analysis and densitometric quantification of total and phosphorylated mTORC1 Ser 2448, p70S6K Thr 389, eEF2 Thr 56, eIFG Ser 1108 in C2C12 myotubes transduced with Ad-CMV-shGFP or Ad-CMV-shGFP-r/mBCKDHA for 48 hr followed by stimulation with 100nM insulin for 15 mins. Statistical analysis was performed using a two-way ANOVA followed by a Tukey’s multiple comparison test; *p <0,05, **p < 0.01, **** p <0.0001 as indicated; *within groups, #between groups. d) Graphical abstract. BCAA catabolic enzyme expression in the gastrocnemius muscles is downregulated with progressive DIO increasing circulating BCKAs (a). Exogenous supply of BCKAs results in impairment of insulin induced AKT phosphorylation by blunting insulin induced PP2A inactivation (b) and decreasing IRS1 phosphorylation. Increased accumulation of intracellular BCKAs by adenoviral overexpression of BCKDK or silencing BCKDH also impairs insulin-induced AKT phosphorylation (c) by activating mTORC1 (d) also resulting in activation of key signaling components of the protein translation machinery. Alternatively, genetic and pharmacological activation of BCKDHA reduces intracellular BCKAs and sensitizes cells to insulin signaling (e) and function (f). Finally, BT2 treatment increases FAO and reduces incomplete FAO (g) further explaining its insulin-sensitizing effects.

Changes in mitochondrial function alter protein translation. As reported in prior studies, ROS overproduction attenuates translation elongation and pauses translation (57, 58). Directing TCA cycle intermediates away from ETC towards translation and lipid and nucleotide biosynthesis suppress mitochondrial respiration. Since mTOR regulates mitochondrial oxygen consumption and oxidative capacity (59), we questioned if BCKAs induced mTORC1 activation in C2C12 cells impacts oxidative and glycolytic metabolism and mitochondrial respiration. Ketoleucine and ketoisoleucine suppressed maximal oxygen consumption rate with no changes in basal respiration (Fig S6a). ATP production was significantly reduced by ketoleucine but remained altered by ketoisoleucine or ketovaline (Fig S6b). Spare respiratory capacity was unchanged following incubation with individual BCKAs (Fig S6c). Our findings are consistent with prior studies demonstrating that BCKAs suppress respiration in cardiac mitochondria (11). Since mitochondrial oxygen consumption and ATP production were altered, we next examined whether ketoleucine treatment influences mitochondrial fatty acid oxidation. Although acute treatment with ketoleucine did not alter ^14^C palmitate oxidation (Fig S6d), CO_2_ derived from the acid-soluble metabolite fraction (incomplete fatty acid oxidation) was increased in cells incubated with ketoleucine (Fig S6e). Together, our findings suggest that BCKA mediated impairment in insulin signaling is associated with increased mTOR signaling and suppressed mitochondrial respiration and fat oxidation.

## 3. Discussion

Prior studies have either supported (17,25,60,61) or challenged (25,62–64) the paradigm that increased circulating BCAAs is a hallmark of IR, T2D and obesity by utilizing genetic (25,26,65) and diet-induced rodent models of obesity and IR. The inconsistencies across different studies relating circulating BCAAs levels to murine obesity and IR and its effects on metabolism are partly attributed to differential dietary intake, tissue uptake, and nutritional status. Coordinate and inter tissue regulation of BCAA metabolizing enzymes also contribute to the reported inconsistencies since there are contradictory findings on the impact of IR on BCAA catabolic enzyme expression in the skeletal muscle (26,32,66). Remarkably, elevations in circulating BCKAs correlated better with the severity of IR and T2D pathogenesis (34, 35). Elevated BCKA is an outcome of impaired action of BCAA catabolizing enzymes, BCKDHA (67, 68), PPM1K (69–71) and KLF15, the transcriptional activator of BCAA metabolism (11, 25). BCKA is generated not only systemically but also intracellularly in tissue capable of BCAA intake and/or BCAA oxidation. A significant amount of BCKA is generated within the skeletal muscle for transport to other organs, like the liver and kidney (72). Moreover, since skeletal muscle is the major organ of insulin-induced glucose disposal (24), it is plausible that loss of BCAA catabolizing enzyme function can overwhelm skeletal muscle BCKA concentration and perturb downstream metabolism. We hypothesized whether increasing BCKAs or targeting BCAA catabolic enzymes exert effects on insulin signaling and function to govern myocyte metabolism.

In this study, we demonstrated tissue specific mRNA and post-translational changes in BCAA catabolizing enzyme in mice subjected to acute (physiological fasting), and chronic (DIO) nutritional stress, an effect that is likely attributed to altered insulin action or lipid content. Exposure to a lipotoxic milieu suppressed BCAA catabolic enzyme expression both *in vivo* and *in vitro* indicating that that fatty acid-induced defects in BCAA catabolism (either BCAA metabolic enzyme inhibition or accumulation of downstream BCAA metabolites) disrupting insulin function. Further, we report that acute exposure of supraphysiological concentrations of individual BCKAs was sufficient to induce insulin signaling impairment in both cardiac and skeletal muscle cells. Inhibiting PP2A, a phosphatase that dephosphorylates AKT, reversed BCKAs effect on insulin signaling. We also demonstrate that reducing intracellular BCKAs by genetic or pharmacological modulation of BCKDH, sensitized cardiac and skeletal muscle cells to insulin. Alternatively, we observed that augmenting intracellular BCKAs by adenoviral silencing of BCKDH or exogenous application of BCKAs in the skeletal muscle cells resulted in increased mTORC1 signaling, a negative regulator of insulin action.

Intracellular BCAAs catabolism is an essential determinant of circulating and intracellular BCKAs levels. Indeed BCAAs catabolic enzymes, BCATm, PPM1K and BCKDHA are downregulated in adipose tissues (70,73–75) of rodents with obesity and IR. In our model of DIO, we observed a significant decrease in downstream genes of BCAAs catabolism, namely, Ivd2 and Mut mRNA levels in the gastrocnemius muscle at 13 wk post-HFHS diet feeding. Similar effects on BCAA catabolic enzyme mRNA expression were made in C2C12 myotubes exposed to incremental concentrations of palmitate. Intriguingly in both the *in vivo* and *in vitro* model of lipid overload, we found a compensatory increase in Hmgcs1 mRNA levels, likely a mechanism to replenish acetyl CoA pools. Our findings are in agreement with a prior study demonstrating decreased expression of BCAAs oxidative genes in human and rodent muscles with IR (32). Moreover, we found an increase in inhibitory phosphorylated BCKDH Ser 293 levels in the gastrocnemius muscle fed HFHS diet for 13 wk likely signifying inhibited BCKDH activity and thereby BCAAs catabolism. Our data are supported by studies in rodent models of obesity demonstrating loss of BCKDH activity and BCAA oxidation in multiple tissues, like adipose, liver and heart (67, 68). In contrast, several groups have also reported that skeletal muscle BCAA catabolic genes are unaltered in diet-induced obese mice (21), ob/ob mice (26) or in IR women with normoglycemia (66). Furthermore, a prior study reported no significant differences in the inhibitory phosphorylation of BCKDHA in the gastrocnemius muscles of mice fed 60% HFD for 10 weeks (21). We attribute this inconsistency to the difference in diet composition and the duration post-feeding and also the nutritional state of the animal before euthanasia. Importantly, our study, for the first time, reports the temporal effect of HFHS feeding on BCAA catabolic enzymes in skeletal and cardiac muscle. Together, our data are consistent with previous reports linking lipid metabolism to remodeling of muscle BCAA catabolism (2,32,76). Strikingly, in our study, acute modulation of nutritional status had a profound and distinct impact on post-translational modification of BCAA metabolizing enzymes in the heart but not skeletal muscle, whereas, chronically, in a setting of DIO, BCAA metabolizing enzymes were altered in the skeletal muscle but not the heart.

We and others have demonstrated that despite unchanged plasma BCAAs, systemic and tissue-specific BCKAs were elevated in obese animals (26) as well as in patients with obesity undergoing cardiac surgery (unpublished data). In our study, we observed marginal increases in circulating BCKAs at 13 wk following HFHS feeding. Emerging studies have shown that BCKAs alter carbohydrate metabolism in multiple insulin-responsive tissues (77),(78). BCKAs inhibit glucose uptake in cardiomyocytes (27) and skeletal muscle (79) and reducing BCKAs levels by cardiac-specific BCATm deletion has been attributed to improvement in basal and insulin-induced glucose oxidation along with increased ATP production (78). Moreover, in the liver and cultured hepatic cells, BCKAs has been demonstrated to upregulate the glycogenolytic enzyme, glycogen phosphorylase (77). However, it was unclear if BCKAs’ effects on metabolism involved changes in insulin signaling. Moreover, a recent study demonstrated that BCAA restriction did not affect cardiac insulin signaling in Zucker fatty rats (80) indicating that downstream metabolites of BCAAs, such as BCKAs, are more potent mediators of insulin signaling. Interestingly despite elevated levels of BCAAs, BCAT2 KO mouse unexpectedly develops resistance to diet-induced obesity and presented with increased glucose disposal, improved glucose tolerance and enhanced insulin sensitivity (81) likely attributed to reduced BCKA levels. We demonstrated that supraphysiological levels of exogenously supplied BCKAs impaired insulin signaling in the skeletal and cardiac muscle cells. Moreover, BCKAs mediated impairment of insulin induced AKT phosphorylation was reversed by okadaic acid, a bonafide inhibitor of PP2A phosphatase that inhibits AKT phosphorylation. Our data are consistent with a recent study in cultured 3T3-L1 adipocytes depicting that BCKAs directly impaired insulin signaling (26).

Prior studies have examined the direct effects of BCAA and BCKA but not a consequence of disrupted BCAA catabolic enzyme expression on cellular metabolism and insulin function. This study, for the first time, demonstrates that altering intracellular BCKAs levels by overexpressing or silencing BCKDHA or BCKDK impacts insulin signaling both in the skeletal and cardiac muscle cells. Overexpressing BCKDHA marginally decreased intracellular BCKAs levels which were sufficient to reverse the inhibition of AKT phosphorylation by direct incubation with BCKAs. Conversely, adenoviral silencing of BCKDHA or overexpressing BCKDK, the kinase inactivating BCKDHA, impaired AKT phosphorylation. Our finding supports the correlation between intracellular BCKAs levels and insulin mediated AKT phosphorylation in the C2C12 myotubes. Similar effects on insulin-mediated AKT phosphorylation was observed in the muscle by incubating with BCKDK inhibitor, BT2 which increased intracellular BCKAs clearance. Our *ex vivo* findings are consistent with the in vivo study wherein BT2 treatment in ob/ob mice improved insulin-induced AKT phosphorylation in the skeletal muscle but not in the liver or white adipose tissue (26). Interestingly, changes in BCKDHA content profoundly affected AKT2 Ser 474 phosphorylation than on AKT1 Ser 473. Previous studies have reported the importance of AKT2 Ser 474 phosphorylation for maximal activation of AKT and insulin-regulated processes (82). Strikingly, BT2 treatment in C2C12 cells with BCKDHA silencing rescued insulin induced AKT phosphorylation on Ser473 but not Ser 474. It is plausible that distinct subunits of BCKDHA exhibit effects on regulating specific AKT isoforms. Nevertheless, our data indicate that lowering intracellular BCKAs levels augment insulin signaling in the muscle. Upon a reduction in intracellular BCKAs levels by either BT2 treatment or BCKDHA overexpression sensitized insulin action as reflected by increased PDH activity. Indeed, hearts with PPM1K deletion (20) and plasmodium with loss of BCKDH function (83) exhibited reduced PDH activity leading to impaired glucose oxidation (84). In our study, increased PDH activity in BT2 treated C2C12 cells corresponded with the upregulation of *Glut1* mRNA expression. BCKAs levels are downregulated in Glut1 overexpressing cardiomyocytes (85) suggesting that BCKA generation and utilization are intricately coupled to glucose metabolism via its effects on insulin signaling.

Muscle insulin signaling is inhibited by mTOR, P70S6K and ribosomal S6, proteins activated by BCAAs (86). A key function of nutrient sensor mTOR is to maintain the available amino acid pool by regulating protein translation. Indeed during DIO, glucolipotoxicity exacerbates IR with concomitant mTOR activation in the heart and skeletal muscle (87, 88). We demonstrate that the three BCKAs individually activated mTOR, with ketoleucine and ketoisoleucine exhibiting pronounced effects which are in agreement with prior studies in cultured adipocytes (26) showing increased mTORC1 activity and defective insulin pathway in response to BCKA exposure (89). mTOR activation increases protein translation and synthesis (90). A concerted action of eukaryotic elongation factor 2 (eEF2), eEF2 kinase (eEF2K) and eukaryotic initiation factor 4G (eIF4G) in conjunction with mTORC1 activation governs translation signaling (55, 56). Depleting C2C12 cells of BCKDHA or treating with BCKAs upregulated eIF4G phosphorylation at Ser 1108, increased phosphorylated mTOR, P70S6K and decreased eEF2 Thr 56 phosphorylation, events which promote translation. We theorize that intracellular BCKAs is an anabolic sensor facilitating mTOR activation and also acting as an endogenous inhibitor of insulin signaling. Interestingly, cardiomyocytes incubated with BCKAs acutely downregulated AKT signaling triggering cell death (91), suggesting that excessive mitogenic signaling including translation, can be an energetically expensive process compromising cellular survival. Supporting this paradigm we and others demonstrate that BCKAs suppress mitochondrial oxygen consumption in C2C12, cardiac (11), neuronal (37) and hepatic cells (77). BCKAs also increased the accumulation of acid soluble metabolites, a signature of incomplete fatty acid oxidation in C2C12 cells.

Taken together, we report that muscle BCAA catabolizing enzymes expression is modulated by acute and chronic changes in insulin during physiological fasting and diet-induced obesity, as well as exposure to increasing concentration of fatty acids. In an environment of chronic lipid overload downregulation of BCAA catabolic enzyme expression accumulates BCKAs causing IR. Cardiac and skeletal muscle cells depleted of BCKDHA and treatment with exogenous BCKAs displayed defective insulin signaling with concomitant activation of mTORC1 and protein translation. BCKAs induced incomplete oxidation of fatty acids and suppressed mitochondrial respiration. Additional studies are warranted to clarify if the metabolic effects of BCKA are a cause or effect of changes in insulin signaling or mitochondria function or protein translation. Moreover, augmenting BCKAs clearance by overexpressing BCKDHA or pharmacological targeting of BCKDK enhanced insulin signaling and glucose utilization. Our findings indicate that BCKAs can independently impair insulin signaling by modulating upstream effectors of AKT (Fig 8d). This study advances the knowledge on the molecular nexus of BCAA metabolism and signaling with cellular insulin action and respiration.

## 4. Materials and methods

### 4.1 Cell lines and culture conditions

C2C12 cells were cultured in DMEM with 10% fetal bovine serum (FBS, Thermo Fisher Scientific). Differentiation was induced with 0.2% fetal bovine serum for 4 to 6 d after cells reached 80% confluence. L6 cells were grown and maintained in α-minimal essential medium (α-MEM; Corning) containing 10% FBS. Differentiation was induced with 2% FBS for 4 to 6d. Cells were grown to 70–80% confluence. All the cell lines were maintained at 37°C in a humidified atmosphere of 5% CO_2_.

#### Primary cardiomyocyte culture

Neonatal rat ventricular cardiomyocytes (NRCMs) were isolated from 2 d old Sprague–Dawley rat pups as described previously (88). Briefly, hearts were excised and the ventricles were minced into small pieces and digested in several steps using collagenase-type 2 (2% W/V; Worthington Biochemical Corporation), DNase (0.5% W/V; Worthington Biochemical Corporation) and trypsin (2% W/V; Worthington Biochemical Corporation) with gentle stirring to dissociate heart pieces into single cells. Following digestion, non-muscle cells and fibroblasts were eliminated using differential plating of the cell suspension for 2 h. Supernatant from differential plating, containing cardiomyocytes was collected and suspended in DMEM/F12 HAM (Sigma) growth medium [containing 10% FBS, 10 μmol cytosine-β-D-arabinofuranoside (ARAC; Sigma), Insulin–Transferrin–Selenium (ITS; Corning), 1X antimycotic/antibiotic solution (Sigma)] and plated in primaria cell culture plates (Corning). The following day, cells were washed and cultured in serum-free DMEM no glucose medium (GIBCO) [containing 10 μmol ARAC, 0.05 mg−1 gentamycin, 1% penicillin–streptomycin and 10 mM glucose] for 24 h.

Adult rat cardiomyocytes (ARCMs) were isolated from adult male Sprague-Dawley rat hearts as described previously (88). Briefly, Langendorff method was used for retrograde perfusion of the isolated heart with Tyrode buffer [containing 1.49 mM KCl, 0.33 mM KH_2_PO_4_, 11.69 mM NaCl, 12.51 mM taurine, 1.206 mM MgSO_4_, 4.766 mM HEPES, 3.604 mM dextrose, and 0.396 mM L-carnitine at pH 7.4] in 5% CO_2_– 95% O_2_ at 37 °C. Following perfusion, the heart was digested with collagenase (60 mg collagenase, 25 μM CaCl_2_ in 75 ml Tyrode buffer) for 30 min in a recirculating mode. The digested heart was excised, and the ventricular pieces finely minced into a homogenous solution. Incremental physiological concentrations of calcium (200 μM, 500 μM and 1 mM) was used to make the ventricular myocytes calcium tolerant, in the presence of 5% CO_2_–95% O_2_. Dead cells were separated from the viable cardiomyocytes by gravity settlement. Cell pellet containing viable cardiomyocytes were resuspended in plating media [containing media 199 (Sigma), 26.2 mM NaHCO3, 25 mM HEPES, 1.24 mM L-carnitine, 137 μM streptomycin (Sigma), 280.6 μM penicillin (Sigma), 10 mM taurine (Sigma) and 1% BSA Fraction V (Roche) at pH 7.4] and seeded at a density of 50–75 × 10^3^ cells/plate on laminin (Roche) coated plates. Plating media was replaced with fresh media after 4 h and cells were treated accordingly.

C2C12 and L6 cells were incubated with serum-free DMEM low glucose, leucine free media (Sigma); NRCMs with serum-free DMEM no glucose media and ARCMs with serum-free media 199 for 16 h before treating with 4-methyl 2-oxopentanoic acid sodium salt (ketoleucine, Sigma), sodium-3-methyl-2-oxobutyrate (ketovaline, Sigma), 3-methyl-2-oxovaleric acid sodium salt (ketoisoleucine, Sigma) or a combination of all three (BCKAs). All the experiments were conducted with 5 mM BCKAs (either individual or a mixture) for 30 mins unless mentioned otherwise. Insulin resistance was induced in C2C12, L6 cells and cardiomyocytes by incubating them in their respective media containing 2% (w/v) fatty acid-free bovine serum albumin (FAF-BSA; Roche) and 0.4 mM sodium palmitate (Sigma) for 16h. Myotubes were incubated with 2% FAF-BSA in the absence of palmitate to mimic an insulin-sensitive state. To examine insulin signaling, cells were incubated with 100 nM insulin (C2C12, L6, ARCMs) or 200 nM insulin (NRCMs) or phosphate-buffered saline (PBS) for 15 min (C2C12, L6) or 20 min (NRCMs, ARCMs). For the BT2 experiments, C2C12 and NRCMs were pre-treated for 20 h with the desired concentration of BT2 (Sigma and Matrix Scientific). Cells were washed once and harvested in ice-cold PBS, followed by centrifugation at 18,000 × g for 10 min at 4°C. Cell pellets were snap-frozen in liquid nitrogen and stored at −80°C until further use.

### 4.2 Fat oxidation

C2C12 myotubes were incubated with either 5mM ketoleucine for 30 mins or 500μM of BT2 for 20 h. Following the treatment, FAO was performed according to a prior study (92). Briefly, serum-starved cells were incubated with 1μCi of ^14^C palmitate (Perkin Elmer) at 37°C for 2 h. Following the incubation, 70% perchloric acid was added to each well and the CO_2_ released was trapped in Whatman paper soaked with 3N NaOH for 1 h. Incomplete fat oxidation was measured by the ^14^C labeling of acid soluble metabolites in the supernatant collected after centrifugation of the media at 500 g for 10 mins at RT. ^14^C levels were determined using a β-counter (Perkin Elmer). The assay was normalized by protein content.

#### Isolated heart perfusion/Fat oxidation

Hearts were perfused aerobically in working mode with Krebs-Henseleit buffer containing 1.2 mmol/litre palmitate prebound to 3% delipidated bovine serum albumin, and 5 mmol/litre glucose, as described previously (21). Mice were euthanized in the fed state, and hearts were dissected and subsequently perfused. Preload and afterload pressure were set to 11.5 and 50mm Hg, respectively, unless otherwise stated. For measurement of fatty acid oxidation rates, hearts were perfused for 30 min with buffer containing 1.2 mmol/litre [9,10-^3^H] palmitate in the presence or absence of 5 mM leucine. Following perfusion, atria were removed, and ventricles were snap-frozen in liquid nitrogen and stored at −80°C until further processing.

### 4.3 Animal Studies

C57BL6J mice were procured from Jackson Laboratory (Bar Harbor, ME, USA). 10 wk old male C57BL/6J mice were divided into 3 groups and either fed ad libitum or fasted for 16 h or refed for 4 h following fasting (n=5 each group). Body weight and blood glucose were measured before euthanasia and liver and ventricle weight was measured after dissection. Gastrocnemius muscles and heart tissue were snap-frozen in liquid nitrogen and stored at −80°C until further processing.

8 wk old C57BL6J male mice were fed either chow or high-fat high sucrose (HFHS) diet and euthanized following overnight fast at 2, 4, 8 and 13 wk (n=5 each group). Diet composition details are included in Table S2. Serum was collected after centrifuging the blood at 5000 rpm for 5 mins. Gastrocnemius muscle and heart tissue were snap-frozen in liquid nitrogen and stored at −80°C until further processing. All protocols involving mice were approved by the Dalhousie University Institutional Animal Care and Use Committee.

### 4.4 Tissue and cell lysate processing and immunoblotting

Frozen hearts and gastrocnemius muscle compartment from mice were powdered and homogenized using a tissue homogenizer (Omni TH, Omni International) in ice-cold lysis buffer [containing 20 mM Tris-HCl, pH 7.4, 5 mM EDTA, 10 mM Na_4_P_2_O_7_ (Calbiochem), 100 mM NaF, 1% Nonidet P-40, 2 mM Na_3_VO_4_, protease inhibitor (10 μl/ml; Sigma) and phosphatase inhibitor (10 μl/ml, Calbiochem)]. Homogenate was centrifuged at 1200 g for 30 min and the supernatant collected for determining protein concentrations. Cell pellets were sonicated in ice-cold lysis buffer and centrifuged at 16,000 g for 15 min. Protein concentrations of the cell and tissue lysates were determined using BCA protein assay kit (Pierce, Thermo Fisher Scientific). Lysates were subjected to SDS-PAGE and proteins were transferred onto a nitrocellulose membrane (Biorad). Proteins were visualized using a reversible protein stain (Memcode, Pierce, Thermo Fisher Scientific) and membranes were incubated with the following primary antibodies listed in Table S3. Immunoblots were developed using the Western Lightning Plus-ECL enhanced chemiluminescence substrate (Perkin Elmer). Densitometric analysis was performed using Image lab software (Bio-Rad) and the quantifications were normalized by total protein loading using Graph pad software (Clarivate).

### 4.5 qPCR analysis

mRNA levels of BCAA metabolizing enzyme and glucose transporter (Gluts) related gene expression were determined in tissues and cells using qPCR by employing validated optimal reference gene pairs as previously described (93). Primer information of the target and reference genes are provided in Table S4. Powdered tissue and cell pellets were homogenized in Ribozol (Amresco)). RNA was isolated as per the manufacturer’s instructions, and RNA quality and quantity were examined using a QIAxcel Advanced System (Qiagen). cDNA was synthesized from 1 μg of RNA using qScript cDNA supermix (Quanta Biosciences) and cDNA samples were stored at −30 °C until further use. qPCR analysis was performed in 96-well plates using PerfeCTa SYBR green Supermix Low ROX (Thermo Fisher Scientific) and a ViiA7 real-time PCR machine (Thermo Fisher Scientific).

### 4.6 AKT activity assay

AKT activity was measured using the KinaseSTAR AKT activity assay kit (Biovision) as per the manufacturer’s instructions. Briefly, C2C12 cells were plated at a density of 2 x 10^6^ and treated with either 100 nM insulin alone for 15 mins or with 5 mM ketoleucine, ketovaline and ketoisoleucine for 30 mins followed by 100 nM insulin for 15 mins. The cells were lysed using cold kinase extraction buffer and pelleted at 13,000 rpm for 10 mins at 4°C. One part of the cell lysate was used for assaying protein concentration. 200µg of the cell lysate was incubated with 2µl of AKT antibody and AKT was immunoprecipitated using Protein A-sepharose beads. Kinase assay was performed by incubating the AKT protein-bound beads with 2µl GSK 3α/ATP mixture at 30°C for 4h. The beads were spun down and supernatant collected and boiled in 2X SDS-PAGE loading buffer separated by 12% SDS-PAGE and analyzed by Western blotting against phosphorylated GSK 3α Ser 21.

### 4.7 Sandwich ELISA assay

AKT phosphorylated at Ser 473 was measured by sandwich ELISA assay using the DuoSet IC ELISA (R&D systems) as per the manufacturer’s protocol. Briefly, C2C12 cells were transduced with either Ad-CMV-shGFP or Ad-CMV-GFP-rm-shBCKDHA (MOI 200) for 48 h or treated with DMSO or 500µM BT2 for 20 h, followed by 100nM insulin treatment for 15 mins. Cells were then solubilized in lysis buffer (1 mM EDTA, 0.5% Triton X-100, 5 mM NaF, 6 M Urea, 1 mM activated Na_3_VO_4_, 2.5 mM Na_4_P_2_O_7_, 10 μg/mL Leupeptin, 10 μg/mL Pepstatin, 100 μM PMSF, 3.0 μg/mL Aprotinin in PBS, pH 7.2-7.4) at a concentration of 1 x 10^7^ cells/ml and centrifuged at 2000 g for 5 min. One day prior to the assay, a 96 well high binding polystyrene plate (Greiner Bio-One) was coated with 6 µg/ml capture antibody (phosphorylated AKT Ser 473) diluted in 1X PBS overnight. For the assay, the supernatant and a seven-point standard curve (100-10000 pg/ml) was diluted 6-fold with diluent 8 (1 mM EDTA, 0.5% Triton X-100, 5 mM NaF in PBS, pH 7.2-7.4) and further diluted in diluent 3 (1 mM EDTA, 0.5% Triton X-100, 5 mM NaF, 1 M Urea in PBS, pH 7.2-7.4). On the day of the assay, the plate was washed thrice with the wash buffer (0.05% Tween® 20 in PBS, pH 7.2-7.4) and blocked at RT for 2 h with blocking buffer (1% BSA, 0.05% NaN_3_ in PBS, pH 7.2-7.4). 100 µl of the sample or standard was added per well and incubated at RT for 2 h followed by aspiration and washing. The samples and standards were then incubated with detection antibody (phosphorylated AKT Ser 473) conjugated with Streptavidin-HRP A at RT for 2 h. Colorimetric reagents were then added, and absorbance measured immediately at 540 nm. The assay was normalized with protein concentration.

### 4.8 PP2A Immunoprecipitation phosphatase assay

C2C12 cells were treated with either 100 nM insulin for 15 min, or pretreated with 250 nM okadaic acid for 45 mins followed by 100 nM insulin for 15 min, or 5 mM ketoleucine for 30 mins followed by 100 nM insulin for 15 mins, or pretreated with 250 nM okadaic acid for 45 mins followed by 5 mM ketoleucine and 100 nM insulin for 15 mins. Cells were lysed with lysis buffer (20 mM imidazole-HCl, 2 mM EDTA, 2 mM EGTA, pH 7.0 with 10 mg/mL each of aprotinin, leupeptin, pepstatin, 1 mM benzamidine and 1 mM PMSF) and 300 µg of lysate was used to measure PP2A activity. Tissue was homogenized on ice and centrifuged at 12,000 rpm for 10 minutes at 4 °C. Cells were sonicated before centrifuging similarly. The supernatants were used to assay PP2A phosphatase activity by a standard kit (EMD Millipore) according to the manufacturer’s instructions. The intensity of the color reaction was measured at 650 nm on a Bio-rad microplate spectrophotometer.

### 4.9 PDH activity assay

C2C12 myotubes were transduced with Ad-mCherry-hBCKDHA (MOI 120) or Ad-GFP-sh-r/mBCKDHA (MOI 200) and their respective controls for 48 h or treated with 500µM BT2 or DMSO for 20 h and PDH activity was measured using the PDH activity assay kit (BioVision). Briefly, 1×10^6^ cells per condition were lysed with 100 µl ice-cold PDH assay buffer and incubated on ice followed by centrifugation at 10,000 rpm for 5min. 50 µl of the supernatant was used and adjusted with 50 µl PDH assay buffer and the reaction was started with 50 µl of the reaction mixture (PDH assay buffer, PDH developer and PDH substrate) and absorbance was measured immediately at 450nm in a kinetic mode for 60 min at 37°C using the Synergy plate reader. NADH standard curve was used to calculate PDH activity in the samples in mU/ml.

### 4.10 Extracellular flux studies

C2C12 cells were plated at a density of 30,000 cells/per well (XF24 cell culture microplate, Seahorse Biosciences) and differentiated to myotubes for 3 d. Mitostress assay was performed as described in a previous study (92). Briefly, cells were washed with XF Assay media (with 20 mM glucose, 1mM sodium pyruvate and 1mM glutamate, without sodium bicarbonate) and incubated for 1 h at 37°C in a non CO_2_ incubator. 2μM of oligomycin, 1μM of FCCP, and 1μM each of rotenone and antimycin A were injected and oxygen consumption rate (OCR) was measured over 100 min. The assay was normalized to protein content. The analysis was performed using the XFe 2.0.0 software (Seahorse Biosciences).

### 4.11 BCKA measurements

#### Serum BCKA extraction

20 μl of the sample was combined with 120 μl of internal standard (ISTD; 4 μg/ml in H_2_O) containing leucine-d3 (CDN Isotopes), 40 μl of MilliQ water, 60 μl of 4M perchloric acid (VWR) were combined and vortexed. Proteins were precipitated in two sequential steps followed by centrifugation at 13,000 rpm for 15 mins at 4°C. Supernatant collected from both steps were combined for measuring BCKAs.

#### Intracellular BCKA extraction

500,000 cells, 120 uL of internal standard (ISTD; 4 ug/ml in H_2_O containing leucine-d3 (CDN Isotopes, D-1973) and 0.8 ng/uL in H_2_O containing sodium-2-Keto-3-methyl-d3-butyrate-3,4,4,4d4 (KIVd7; CDN Isotopes), 120 µl of 6M perchloric acid (VWR) were combined and homogenized with a tissue homogenizer. Proteins were precipitated in two sequential steps, followed by centrifugation at 16,500 g for 15 mins at 4°C. Supernatant collected from both steps were combined and split into two portions for measuring BCKAs. Samples were derivatized according to previously established protocols (94, 95).

#### BCKAs Derivatization and Quantification

150 µL of extract and 500 µL of 25 mM OPD in 2M HCl (made from o-Phenylenediamine, 98%; VWR) were combined. The mixture was vortexed and then incubated at 80°C for 20 min followed by incubation on ice for 10 min. The derivatized extract was centrifuged at 500 g for 15 min and the supernatant transferred to a tube containing 0.08 g sodium sulfate (VWR) and 500 µl of ethyl acetate (ethyl acetate; VWR) following which they were centrifuged at 500g at RT for 15 min. This step was repeated twice and the supernatant collected were vacuum centrifuged at 30°C for 45 min Samples were then reconstituted in 64 µL of 200 mM ammonium acetate (made from; ammonium Acetate, 98%; VWR) and transferred to amber glass UPLC vials (Waters). BCKAs were quantified with a Waters Acquity UPLC, Xevo-μTandem Mass Spectrometer and an Acquity UPLC BEH C18 (1.7 µm, 2.1mm X 50 mm; Waters) and ACQBEHC18 VanGuard (130Å, 1.7µm, 2.1mm X 5mm; Waters) using multiple reaction monitoring (MRM) and internal standard calibration as per a prior study (96).

## Declaration

### Funding & Acknowledgement

This work was supported by Natural Sciences and Engineering Research Council of Canada (RGPIN-2014-03687), Diabetes Canada (NOD_OG-3-15-5037-TP & NOD_SC-5-16-5054-TP) and New Brunswick Health Research Foundation grants to T.P.; D.B., was funded by postdoctoral fellowships from New Brunswick Health Research Foundation and Dalhousie Medicine New Brunswick.

### Conflict of Interest

The authors declare no conflict of interest.

### Availability of data and materials

All data and materials used in the current study are available from the corresponding author upon request.

### Consent for publication

Not applicable

